# Production of cellobiose from ionic liquid-treated cellulose using the highly thermostable cellobiohydrolase HmCel6A-3SNP at 80°C and analysis of enzymatic accessibility to the substrate

**DOI:** 10.64898/2026.05.30.728921

**Authors:** Takeshi Ara, Tsutomu Kodaki, Yoichi Ogawa, Tomoya Imai, Seiji Takahashi, Yoshitsugu Hirose, Daisuke Shibata, Toshiyuki Nohira

## Abstract

Cellobiose is an important disaccharide used in food, health, and biorefinery applications, but its efficient enzymatic production from crystalline cellulose remains challenging. In this study, crystalline cellulose was dissolved in ionic liquids and regenerated by dilution, and subsequently hydrolyzed at 80°C using a highly thermostable cellobiohydrolase, HmCel6A-3SNP. The enzyme retained activity in the presence of low concentrations of ionic liquids. Among the pretreatment conditions tested, cellulose treated with 1-butyl-3-methylimidazolium chloride ([Bmim]Cl) showed the highest enzymatic digestibility. After washing to remove residual ionic liquid, the reaction produced reducing sugars at levels 1.5-fold higher than those obtained in the presence of 10% [Bmim]Cl, with cellobiose accounting for approximately 96% of the products. Under the optimized conditions, the hydrolysis yield reached ∼36% after 48 hr. Structural analyses using birefringence imaging, electron microscopy, and Fourier transform infrared spectroscopy indicated that higher-order structural changes in regenerated cellulose strongly influence enzymatic accessibility. These results demonstrate the potential of combining ionic-liquid pretreatment with thermostable enzymes for selective cellobiose production from cellulose.

## Introduction

Cellobiose (4-O-(beta-D-glucopyranosyl)-D-glucopyranose) is a disaccharide composed of two molecules of D-glucose linked by a beta-1,4 bond. Cellobiose is a component of cellulose, which is the most abundant organic substance on Earth. However, the natural abundance of cellobiose is quite low. Therefore, inexpensive, large-scale production of cellobiose using abundant cellulose as a starting material is a promising approach for producing sustainable and functional bio-components. Methods have been developed to convert cellobiose as a starting material into various bioplastic raw materials (Li *et al*. 2014, Qu *et al*. 2016). Chemical modification of cellobiose at specific sites can produce functional materials with polymer stabilization and gelation abilities (Berson *et al*. 2008, Imura *et al*. 2014). Combining cellobiose with other polymers can produce functional biocomposite materials (Isono *et al*. 2020, Sugiura *et al*. 2021). Also, cellobiose is expected to function as a prebiotic when added to food, especially in the health industry, to improve the intestinal environment and maintain digestive health (Paßlack *et al*. 2021).

In industrial cellobiose production, chemical methods such as acid or alkali hydrolysis and organic solvent treatment of cellulose are commonly used. However, these methods inevitably generate by-products that complicate the separation and purification processes. They also require special facilities and equipment for disposing of acid, alkali, and organic solvents, as well as for treating the process under high-temperature and high-pressure conditions. These requirements result in high production costs (Zhang *et al*. 2012, Abolore *et al*. 2024). Conversely, enzymatic treatment of cellulose to produce cellobiose has several advantages, including reduced by-product production, easier purification, production under milder conditions than chemical methods, and easier waste liquid treatment. Several reports have been published on producing cellobiose from cellulose via enzymatic degradation (Homma *et al*. 1993, Pramanik *et al*. 2013, Vanderghem *et al*. 2010).

At least three types of cellulolytic enzymes are involved in cellulose degradation in nature. In the first step, endo-type cellulase cleaves inside the cellulose molecule. Next, exo-type cellulase (cellobiohydrolase, CBH) cleaves cellobiose from the cut end, which is then rapidly degraded to glucose by glucosidase. Industrially used cellulases are often cellulase mixtures produced by filamentous fungi as they are relatively inexpensive. In order to increase cellobiose production efficiency, methods have been proposed to either inhibit cellobiose degradation (Homma *et al*. 1993, Pramanik *et al*. 2013) or to physically separate cellobiose before degradation occurs (Vanderghem *et al*. 2010).

Since these enzymes used for cellobiose production are not heat-resistant, the reactions are usually carried out at 55°C or lower (Haki and Rakshit 2003). In industrial production, the cellulose material is typically impure and contains various microorganisms. This leads to problems such as bacterial proliferation during cellulose decomposition and impurity formation. Furthermore, as the decomposition proceeds, the viscosity of the reaction liquid increases in line with the rising sugar concentration, requiring considerable energy for stirring the reaction vessel. Enzymatic reactions that occur at high temperatures (80°C or higher) can inhibit microbial growth. For example, thermostable amylases are industrially used to produce glucose from starch and play an important role in improving the efficiency of bioethanol production (de Souza and de Oliveira Magalhães. 2010). However, to the best of our knowledge, there are no reports on using thermostable cellulases in the industrial production of cellobiose from cellulose. Pretreatment of cellulose is crucial for efficient enzymatic saccharification.

Traditionally, acid, alkali, and organic solvents have been used for pretreatment. However, it has been pointed out that these methods have a disposal cost problem originated from large amounts of waste liquid generated during pH adjustment (Abolore *et al*. 2024). Therefore, the use of ionic liquids for pretreatment of cellulose has been studied, as they offer advantages such as non-volatility and non-flammability. They are expected to improve safety and reduce the amount of waste liquid (Barbará *et al*. 2025). However, since ionic liquids sometimes inhibit enzyme activity, studies have been conducted into the development of ionic liquid-tolerant cellulases and the washing of cellulose treated with ionic liquids. *Penicillium verruculosum* CBH-I mutant R1 exhibited approximately 70% activity against 5% 1-butyl-3-methylimidazolium chloride ([Bmim]Cl) or 2% choline chloride in a reaction using microcrystalline cellulose Avicel as substrate at 65°C for 1 hr (Pramanik *et al*. 2021). *Talaromyces emersonii* Cel7A mutant 1M10 revealed 60% activity against 20% (w/w) 1-ethyl-3-methylimidazolium acetate at 50°C for 16 hr (Wolski *et al*. 2016). The halophilic archaeon *Halorhabdus utahensis* CBH1 showed 100% activity against 20% (w/w) 1-allyl-3-methylimidazolium chloride ([Amim]Cl) and 60% activity against 30% (w/w) [Amim]Cl at 37°C for 1 hr using carboxymethyl cellulose (CMC) as substrate (Zhang *et al*. 2011). These reports suggested that ionic liquid-tolerant CBH enzymes exist in nature but their thermotolerance is insufficient for industrial usages.

In order to efficiently optimize the pretreatment, it is crucial to understand the structural changes caused by cellulose pretreatment. Despite numerous observations using electron microscopy, polarizing microscopy, nuclear magnetic resonance (NMR) spectroscopy, Fourier transform infrared (FTIR) spectroscopy and molecular dynamics simulations on native cellulose, our knowledge of the structural changes in cellulose during pretreatment remains insufficient (Rabideau 2013, Li *et al*. 2015, Endo *et al*. 2016, Li *et al*. 2017, Li *et al*. 2018, Dissanayake *et al*. 2018, Uto *et al*. 2018, Brehm *et al*. 2019, Rajeev and Basavaraj 2019, Chen *et al*. 2020). Birefringence is a structural characteristic of cellulose that can be evaluated with optical microscopy. Recently, the higher order structure of cellulose was evaluated using a birefringence camera (Uetani *et al*. 2019). However, there is no report on the change in the higher-order structure of cellulose treated with ionic liquids using birefringence image analysis.

In the previous research, we metagenomically isolated a hyper-thermostable cellobiohydrolase (CBH) named HmCel6A from hot spring sediments. We then engineered its variant, HmCel6A-3SNP (Takeda *et al*. 2022). The CBH HmCel6A-3SNP is highly stable and active around 80°C. Here we report the production of cellobiose in the purity of 96% from ionic liquid-treated regenerated cellulose using the CBH at 80°C. We also evaluated the structural accessibility of cellulose prepared with ionic liquids, such as 1-butyl-3-methylimidazolium chloride ([Bmim]Cl) or choline acetate (ChoAc) using a birefringence camera and FTIR spectroscopy. Taken together, these results suggest that the higher-order structural changes in cellulose, caused by ionic liquids, impact the accessibility of enzymes to their substrates.

## Materials and Methods

### Chemicals

Cellulase from *Trichoderma reesei* (*T. reesei*), 1-butyl-3-methylimidazolium chloride ([Bmim]Cl), and Avicel PH-101 (Avicel) were purchased from Merck & Co., Inc., (Rahway, NJ, USA). Cellulose from *Aspergillus niger* (*A. niger*) and choline acetate (ChoAc) were purchased from FUJIFILM Wako Pure Chemical Co., (Tokyo, Japan). Cellulase from *Trichoderma viride* (*T. viride*) was purchased from Cosmo Bio Co., LTD., (Tokyo, Japan).

### Chemical synthesis of codon-optimized CBH HmCel6A-3SNP and construction of expression vectors

The gene sequence encoding the hyper-thermostable cellobiohydrolase (CBH) HmCel6A-3SNP (Takeda *et al*. 2022) was optimized for codon usage with the GeneOptimizer algorithm (Thermo Fisher Scientific, Waltham, MA, U.S.A.). The following sequences were chemically synthesized by Genscript Japan Co., Ltd. (Tokyo, Japan). These sequences include the signal peptide (31 amino acids) of *Nicotiana tabacum* RP-1a (accession no. X12737), HmCel6A-3SNP (accession no. LC163906), the TEV protease recognition site (ENLYFQG), a flexible linker (GGGS)_3_, amino acids from 4 to 93 of *Zea mays* g-ZEIN (accession no. AF371261), a linker and HiBit (VSGWRLFKKIS), and a linker and His_6_ tag (KDEL). These sequences were included to facilitate the efficient purification of the expressed protein. They were ligated sequentially from the N-terminus.

The synthetic sequence was inserted into pUC57, and the HmCel6A-3SNP sequence was amplified by PCR using the following primer sets: pET15b-CBH2-Fw: 5’-CCGCGGCAGCCATCTCGATAACCCATTCATTGGA -3’, and pET15b-CBH2-Rv: 5’-AGCCGGATCCTCGAGTTAGGGCTGAATTGGTGGTA -3’. KOD-plus Neo (TOYOBO CO, LTD., Osaka, Japan) was used for PCR. The HmCel6A-3SNP amplified fragment and the pET-15b expression vector (Merck & Co., Inc., Rahway, NJ, USA) were digested with *Nde* I and *Xho* I (New England Biolabs Japan, Tokyo, Japan) respectively, and were purified using FastGene Gel/PCR Extraction Kit (NIPPON Genetics, Tokyo, Japan). The *Nde* I-*Xho* I fragment of pET-15b was terminally dephosphorylated with Alkaline Phosphatase (*E. coli* C75) (Takara Bio, Shiga, Japan) and ligated with HmCel6A-3SNP using the seamless Ligation Cloning Extract method (Motohashi 2015) to generate pET-CBH2. The DNA sequence of pET-CBH2 was confirmed by sequencing (Azenta Life Sciences, Tokyo, Japan).

### Heterologous expression and purification of the recombinant protein in *Escherichia coli*

*Escherichia coli* JM109 (DE3) was transformed with pET-CBH2 and the resulting transformant was inoculated into 4 mL of LB medium containing 50 μg/mL of ampicillin. The culture was incubated overnight at 37°C and 180 rpm. Then, 0.5 mL of this preculture was inoculated into 200 mL of LB medium containing 50 μg/mL ampicillin and cultured at 37°C and 180 rpm until the OD600 reached 0.4 to 0.5. Isopropyl beta-D-1-thiogalactopyranoside (IPTG) was added to the culture medium to achieve a final concentration of 0.1 mM, and the culture was further incubated at 18°C and 180 rpm for 24 hr. The culture medium was centrifuged at 5,000 ×*g* for 15 min at 4°C. The pellet was resuspended in 5 mL of sterile ultrapure water (SDW), and then the mixture was centrifuged again under the same conditions to wash the cells. The resulting pellet was stored at -30 °C until use.

The *E. coli* pellet was suspended in 5 mL of 20 mM phosphate buffer (pH 7.4) containing 0.5 M NaCl, 20 mM imidazole, 1 mg/mL lysozyme (FUJIFILM Wako Pure Chemical Co., Tokyo, Japan), and either 1×cOmplete (Roche Ltd., Basel, Switzerland) or 1 mM PMSF, and the mixture was incubated on ice for 1 hr. The suspension was homogenized using an ultrasonic crusher (Sonifier 250, Branson, Brookfield, CT, U.S.A.) with the following settings: Output Control 2 and Duty Cycle Constant (10 sec ON, 30 sec OFF on ice) for 5-10 cycles. The solution was centrifuged at 8,000×*g* for 15 min at 4°C. The resulting supernatant was collected in a new tube as a soluble fraction. Then, the pellet was resuspended in 2.5 mL of SDW and collected as an insoluble fraction.

A HisTrap HP 1 mL column (Global Life Sciences Technologies Japan, Tokyo, Japan) was washed with 5 mL of ultrapure water and then equilibrated with 5 mL of 20 mM phosphate buffer (pH 7.4) containing 0.5 M NaCl and 20 mM imidazole. The solution was filtered through a 0.45 μm filter. The His-tag fusion protein was bound by passing the entire soluble fraction through the column, and the flow-through fraction was collected. Subsequently, the column was washed with 10 mL of 20 mM phosphate buffer (pH 7.4) containing 0.5 M NaCl and 20 mM imidazole, and the liquid that passed through the column was collected as a wash fraction. The His-tag fused protein was eluted by passing 5 ml portions of 20 mM phosphate buffer solution (pH 7.4) containing 0.5 M NaCl and imidazole at concentrations of 50, 100, 200, 300 or 500 mM sequentially through the column. The eluates were collected into microtubes in 1 mL increments.

The purity of the collected protein fractions was confirmed by SDS-PAGE. The his-tagged HmCel6A-3SNP fractions were pooled and concentrated using Amicon Ultra-15 (10 kDa) (Merck & Co., Inc., Rahway, NJ, USA) while the buffer was exchanged for 50 mM Tris-HCl buffer (pH 7.5) containing cOmplete EDTA free Protease Inhibitor Cocktail.

### Cellulose pretreatment with ionic liquids

For the ion liquid treatment, 100, 200 or 300 mg of solid [Bmim]Cl or ChoAc was melted at 100 °C for 10 min. Then, 10 mg of Avicel was added to each solution, and the mixture was kept at 100 °C for an additional 30 min. The final ionic liquid concentration was adjusted to 10%, 20%, or 30% (w/v) by adding 50 mM sodium acetate buffer (pH 5.5). Note that cellulose was precipitated as regenerated cellulose in the presence of an aqueous solution within an ionic liquid. The buffer or ethanol washing experiment was performed by adding 0.5 mL of either 50 mM sodium acetate buffer (pH 5.5) or 99% ethanol to the solution of 10 mg of Avicel treated with 100 mg of either [Bmim]Cl or ChoA, as described above. The mixture was kept at 70°C for 30 min and then centrifuged to remove the supernatant. The washing process was repeated twice. The resulting pellet was used as the washed sample, and the unwashed pellet was used as the control. For the ethanol washing, the washed pellet was resuspended in 0.5 mL of 50 mM sodium acetate buffer (pH 5.5) and kept at 70°C for 30 min. The ethanol was removed by centrifugation and this process was repeated twice.

### Enzyme assay

CBH activity was measured using the dinitrosalicylic acid (DNS) method. CBH HmCel6A-3SNP was reacted with phosphoric acid-swollen cellulose (PASC), [Bmim]Cl-or ChoA-treated Avicel at 80°C for various periods of time. PASC was prepared using the previously described method (Wood 1971). After the reaction, the supernatant was collected by centrifugation and used for the hydrolysis assay (Miller 1959). D-glucose was used as the standard. One unit (U) is defined as the amount of enzyme that produces 1 μmol of D-glucose per minute under the conditions specified in the assay procedure. All measurements were performed three times using the same reaction batch, and the mean values are shown. The error bars represent the standard deviation (SD).

### High-performance liquid chromatography

The sugars in the enzyme reaction mixture were identified using high-performance liquid chromatography (HPLC) (Tosoh Corporation, Tosoh Corporation, Tokyo, Japan) equipped with a refractive index detector and an HPX-87H ion-exclusion column (Bio-Rad Laboratories, Hercules, CA, USA). The mobile phase of 5 mM H_2_SO_4_ was used at a flow rate of 0.6 mL/min. All measurements were performed three times and the mean values are shown. Error bars indicate the standard deviation (SD).

### Birefringence imaging of dissolved cellulose

Dissolved crystalline cellulose was observed under a microscope equipped with a birefringence detector. First, 0.2 g of ionic liquid was melted by heating at 100°C for 10 min. Next, 20 mg of Avicel was added, and the mixture was kept at 100°C for 30 min. After the samples were physically mixed using a plastic pipette tip, each mixture was smeared onto a cleaned glass slide and sealed with a cover glass. The glass slides were observed using an optical microscope Axioplan (Carl Zeiss, Oberkochen, Germany) equipped with a birefringence detector PA-micro S (Photonic Lattice Inc., Sendai, Japan).

In addition, the dissolution process of crystalline cellulose was observed using an optical microscope equipped with a temperature-controlled stage in the above setup. First, Avicel was added to the ionic liquid at 75°C for 10 min. Then the sample was physically mixed with the ionic liquid using a plastic pipette tip, and placed between a glass slide and a cover glass. The glass slide was then placed on a microscope glass plate heat plate ThermoPlate TPH-SX-100 glass slide heater (Tokai Hit Co. Ltd., Fujinomiya, Shizuoka, Japan) and observed using an optical microscope Axioplan (Carl Zeiss, Oberkochen, Germany) equipped with a birefringence detector PA-micro S (Photonic Lattice Inc., Sendai, Japan). Once the specified temperature was reached, it was held for 2 min before an image was recorded. Immediately after capturing the image, the temperature was raised by 5°C and maintained for additional 2 min before capturing another image. This process was repeated until the temperature reached 100°C. It takes approximately 10 sec for the detector to record a single image.

### Scanning electron microscope analysis

The cellulose treated with ionic liquid was washed with 100 mM sodium acetate buffer (pH 4.5). The solid portion of the washed cellulose was centrifuged at 20,000 g for 10 min and dehydrated with increasing ethanol concentrations: 50%, 70%, 90%, and 100%. The samples were suspended in a mixture of ethanol and tertiary butyl alcohol and incubated for 15 min. The samples in tertiary butyl alcohol were frozen on ice for several minutes and lyophilized overnight in a lyophilizer. The dried samples were fixed on copper stubs with electronically conductive adhesive (DOTITE, Fujikura Kasei Co. Ltd., Tokyo, Japan) and coated with platinum using an ion sputtering metal coating apparatus JEC-3000 FC (JEOL Inc., Akishima, Japan). The platinum-coated samples were observed using a field emission scanning electron microscope (FE-SEM) JSM 7800 F Prime (JEOL Inc., Akishima, Japan) at an acceleration voltage of 1.5 kV and a working distance of 6 mm.

### FTIR analysis

The cellulose samples treated with ionic liquids were washed with 100 mM sodium acetate buffer (pH 4.5). The lyophilized powder was placed on a diamond probe and single-bond attenuated total reflection (ATR) was measured using an FTIR spectrometer Spectrum One (PerkinElmer Inc., MA, U.S.). The spectrum was acquired by 16 times integration with a resolution of 4 cm^-1^. Measurements were repeated five times for each sample (Avicel, [Bmim]Cl-treated Avicel, ChoAc-treated Avicel, PASC). Background subtraction of the spectral data was performed using the program supplied with the spectrophotometer, and the data was analyzed using the Python package specio (Lemaitre 2017). The following spectral calculations were then processed using Python 3.9: spectra with transmittance higher than 20% were selected, and converted from transmittance to absorbance. The spectra were subjected to Savitzky-Golay spectral smoothing followed by principal component analysis (PCA) using the scikit-learn package in the Python library.

## Results

### Enzyme activity in the presence of ionic liquid

We examined the effects of ionic liquids on the CBH activity in ionic liquid solutions (Fig. 1). [Bmim]Cl or ChoAc was added to 0.1 mL of 1% PASC to achieve final concentrations of 10%, 20% and 30%, respectively. The reaction was initiated by adding

**Figure 1.**
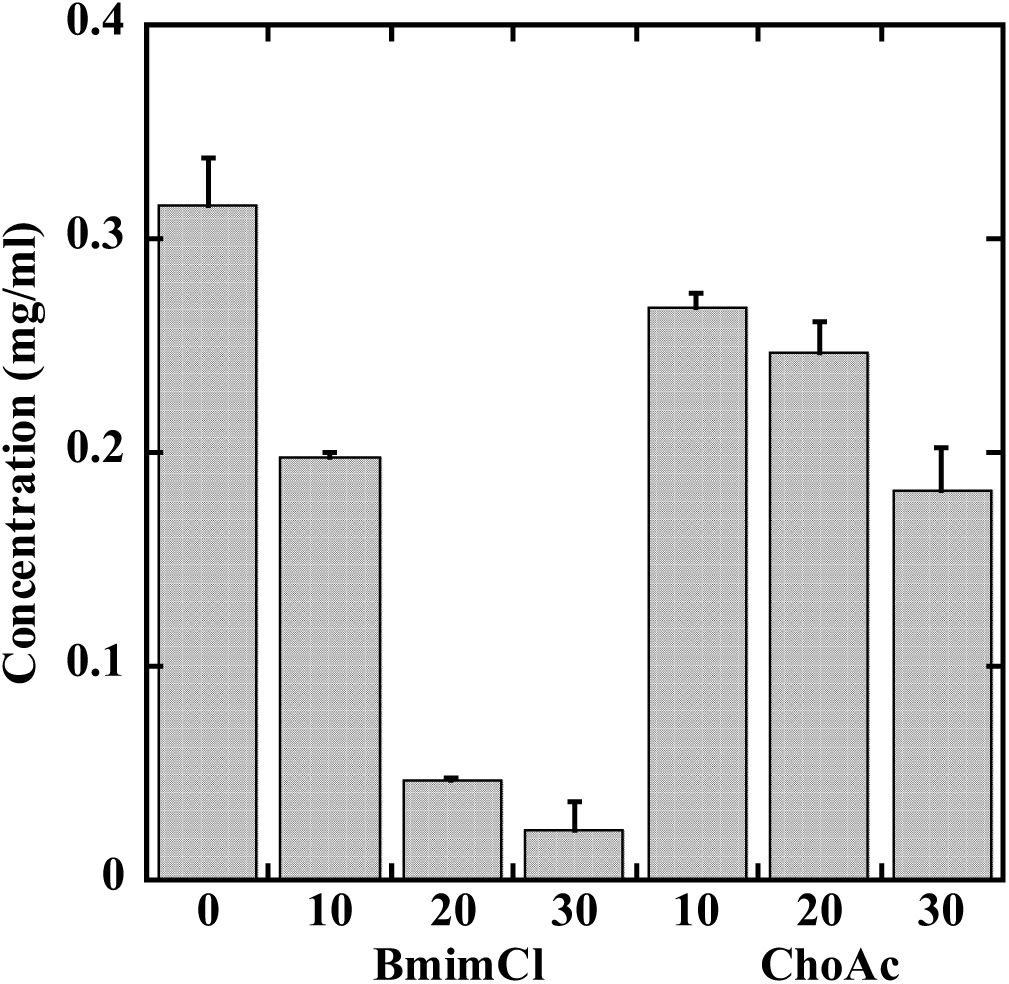
Enzyme activity in the presence of ionic liquid. The enzyme reaction was carried out at 80°C for 1 hr with 1% PASC in the presence of [Bmim]Cl or ChoAc at the respective concentrations (%), and the reducing power of sugars was measured by the DNS method. The error bars in the figure represent the standard deviation (SD).

0.003 U of CBH HmCel6A-3SNP, and the reaction mixture was incubated at 80°C for 1 hr with constant agitation since the substrate is insoluble in the buffer solution. After centrifugation, the reducing power of sugars in the supernatant was measured using the DNS method to quantify the enzymatic saccharification (Fig. 1). The presence of 10%, 20% and 30% [Bmim]Cl decreased the activity to 63%, 15% and 7% of the initial activity, respectively. ChoAc exhibited less inhibitory effects than [Bmim]Cl and retained 85%, 78% and 58% of activity in the presence of 10%, 20% and 30% ChoAc, respectively.

### Time course of sugar formation from regenerated celluloses in diluted ionic liquid solutions

After treatment of the crystalline cellulose Avicel with [Bmim]Cl, 50 mM sodium acetate buffer (pH 5.5) was added to adjust the final ionic liquid concentration to 10% or 20%, respectively. Similarly, the final concentration of ChoAc was adjusted to 10%, 20% or 30%, respectively.

Similar to PASC, the regenerated cellulose formed from ionic liquid treatment became insoluble upon buffer addition. Then, CBH HmCel6A-3SNP (0.038U) was added to 1 mL of 1% cellulose suspension in the presence of various ionic liquid concentrations. The mixture was incubated at 80°C for 25 hr with constant agitation. The enzymatic saccharification was quantified using the DNS method (Fig. 2). Untreated Avicel and the regenerated cellulose PASC were included in the experiment as controls.

**Figure 2.**
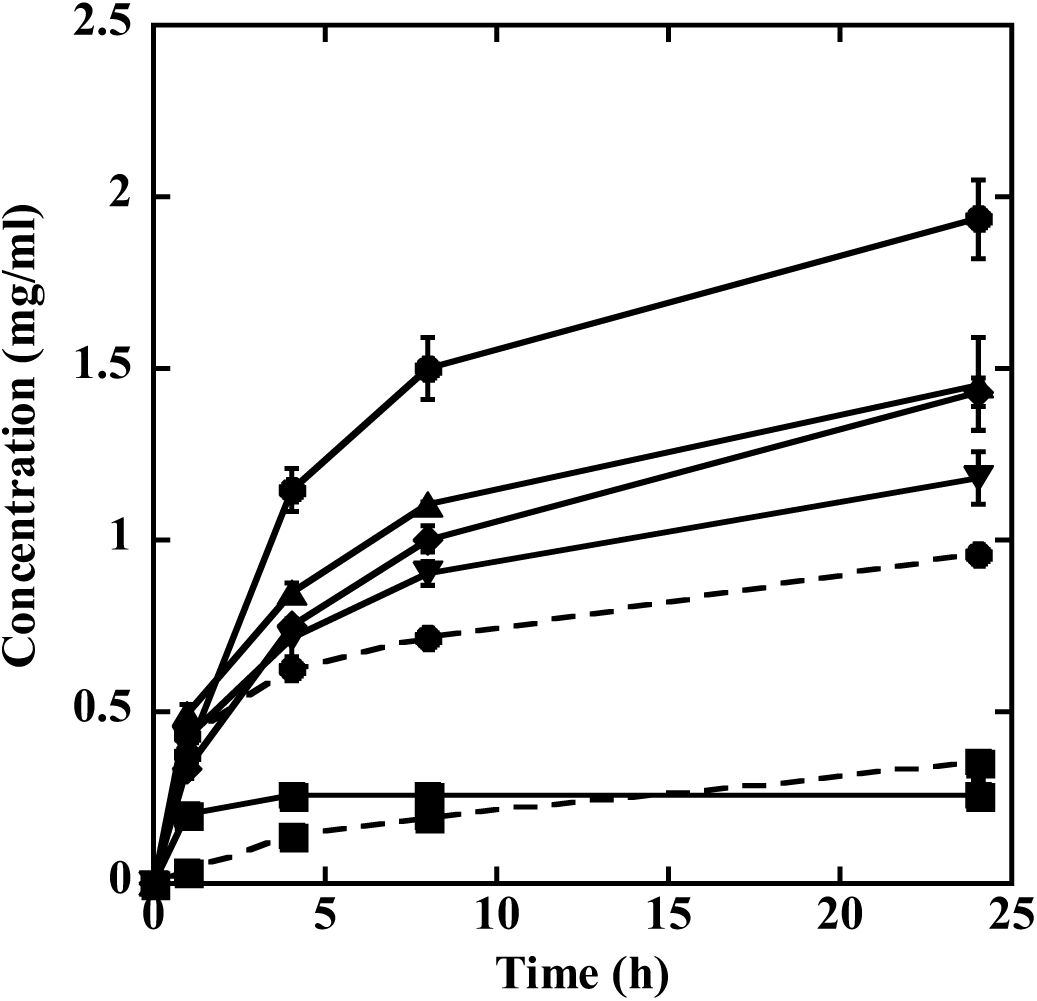
Formation of Sugars from Avicel in Various States over Time. CBH HmCel6A-3SNP (0.038U) was added to 1 mL of a solution (50 mM sodium acetate (pH 5.5) with [Bmim]Cl or ChoAc) containing 10 mg of regenerated Avicel or native Avicel or PASC in each preparation condition. The reaction was carried out at 80 °C for up to 24 hr and the reducing power of the sugar was measured with time by the DNS method. Native Avicel: dashed square; PASC: dashed circle; 10% [Bmim]Cl: solid circle; 20% [Bmim]Cl: solid square; 10% ChoAc: solid diamond; 20% ChoAc: solid triangle; 30% ChoAc: solid inverted triangle. The error bars in the figure represent the standard deviation (SD).

The enzymatic saccharification of regenerated celluloses in the presence of ionic liquids was much higher than that of untreated Avicel except in the presence of 20% [Bmim]Cl. The highest amount of sugar was produced in the presence of 10% [Bmim]Cl. After 24 hr, the 10% [Bmim]Cl mixture produced 5.4 times more sugar than the untreated Avicel and 2.0 times more than PASC. In the presence of ChoAc, more sugar was produced than in the presence of untreated Avicel at any concentration; however, in all cases, it was produced at a lower rate than in the presence of 10% [Bmim]Cl. For PASC, approximately the same amount of sugar was formed at 1 hr as in the presence of 10% [Bmim]Cl, but the rate of formation decreased thereafter.

### Optimization of reaction conditions to increase the sugar production

The regenerated cellulose prepared from Avicel with 10% [Bmim]Cl or 10% ChoAc was washed with 50 mM sodium acetate buffer (pH 5.5) 1, 2, or 3 times to remove excess ionic liquid, and then subjected to the CBH reaction (Fig. 3). The enzymatic activity toward substrates prepared with [Bmim]Cl or ChoAc increased with the buffer washing treatment. The amount of sugars formed by [Bmim]Cl or ChoAc increased by 1.5- or 1.3-fold, respectively, after twice washing with the buffer, compared to the amount without washing. The highest amount of sugars was obtained by using [Bmim]Cl with after twice washing. The effect did not change significantly with an increase in the number of washes, thus we decided to treat twice as the standard procedure.

**Figure 3.**
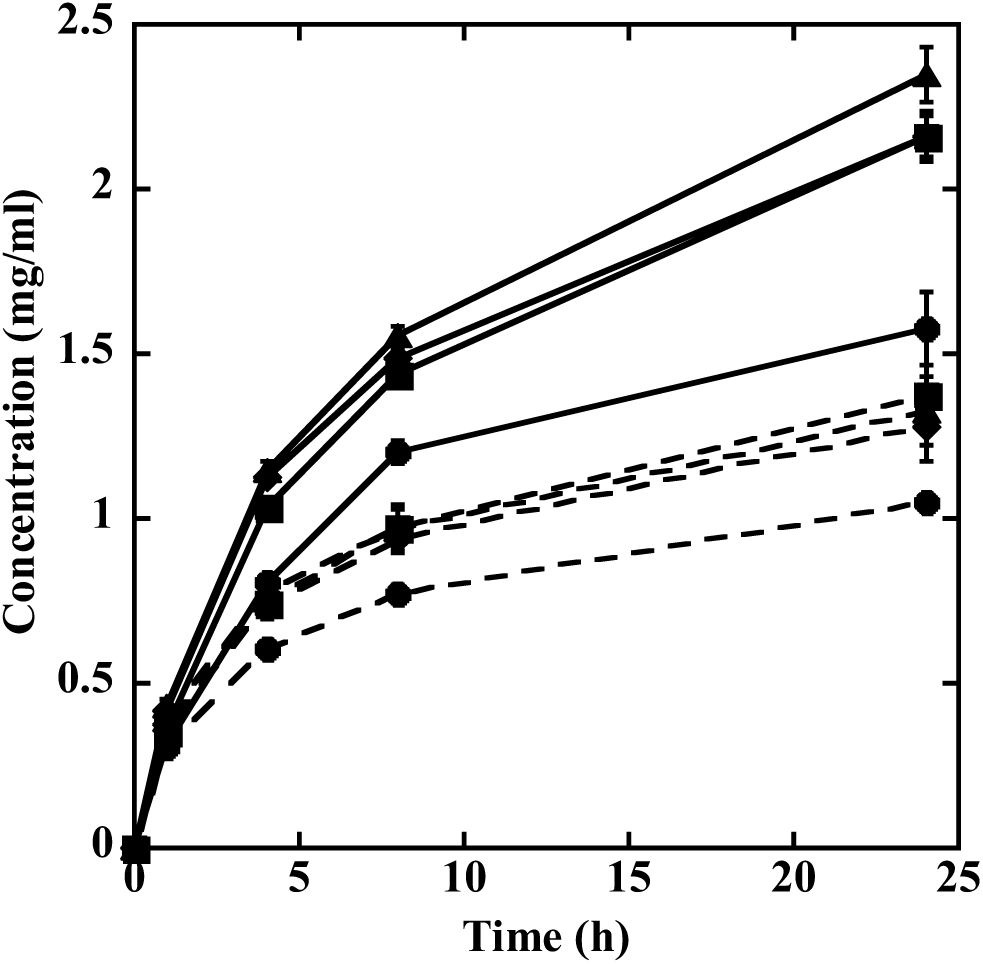
Effect of washing regenerated celluloses with acetate buffer on sugar production. The 10 mg regenerated Avicel treated with 10% [Bmim]Cl or 10% ChoAc was washed 1-3 times with 50 mM sodium acetate buffer (pH 5.5). Then, the resulting pellet was resuspended in 1 mL of 50 mM sodium acetate (pH 5.5) and added 0.038U of CBH HmCel6A-3SNP. The reaction was carried out at 80°C for up to 24 hr and the reducing power of the sugar was measured with time by the DNS method. [Bmim]Cl: solid line; ChoAc: dashed line. 0 time wash: circle, 1 time wash: square, 2 times wash: triangle, 3 times wash: diamond. The error bars in the figure represent the standard deviation (SD).

Ethanol treatment increased both the amount of ionic liquid removed and the amount of sugars produced. Avicel was dissolved in 10% [Bmim]Cl or 10% ChoAc and washed with ethanol 1-3 times. Then, it was washed 2 times with 50 mM sodium acetate buffer (pH 5.5) to remove the ethanol, and added the buffer to make the final volume 1 mL. CBH HmCel6A-3SNP (0.038U) was added to the solution, and the amount of sugars formed over time was measured using the DNS method (Fig. 4). Regardless of the number of ethanol washes, the amount of sugars produced was nearly the same as after the buffer washing alone. Considering the reuse of ionic liquids, they are easier to recover from ethanol than from an aqueous solution. Therefore, the process of ethanol washing may be included in the process when considering the reuse of ionic liquids, that is a critical step for industrial applications (Wojtczuk *et al*. 2020, Ovejero-Pérez *et al*. 2021).

**Figure 4.**
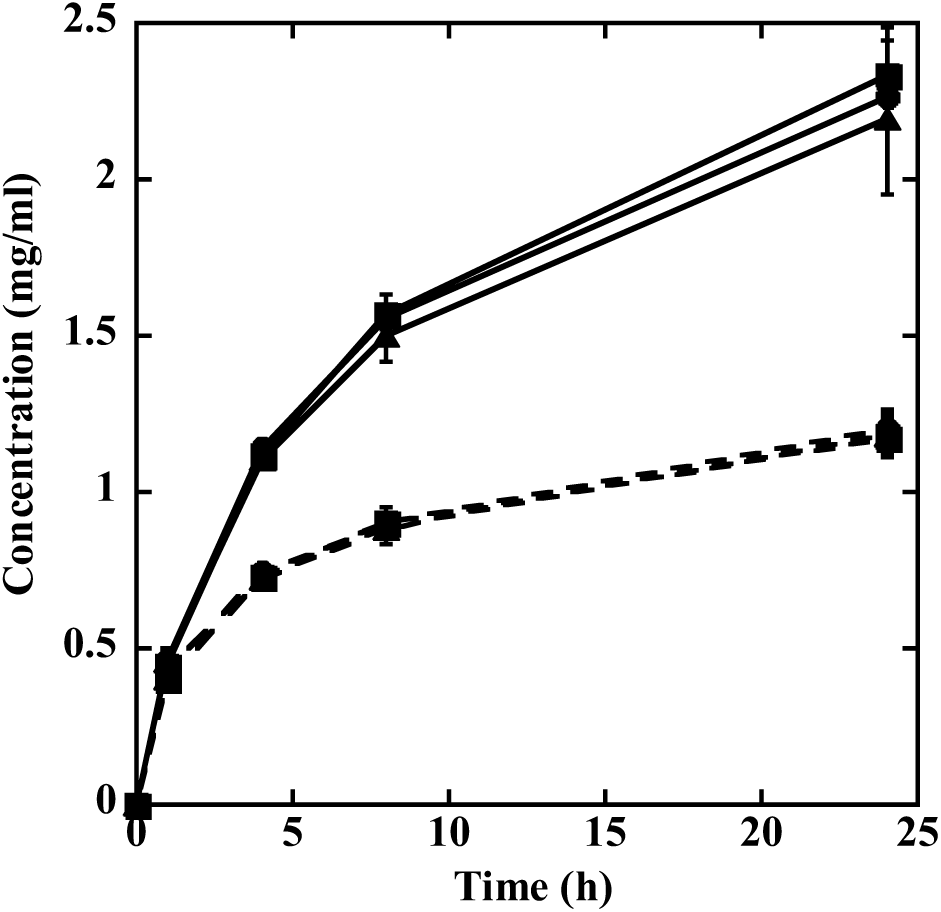
Effect of ethanol treatment. The 10 mg regenerated Avicel treated with 10% [Bmim]Cl or 10% ChoAc was washed 1-3 times with 99% ethanol, and then washed with twice 50 mM sodium acetate buffer (pH 5.5). Finally, the resulting pellet was resuspended in 1 mL of 50 mM sodium acetate (pH 5.5) and added 0.038 U of CBH HmCel6A-3SNP. The reaction was carried out at 80 °C for 24 hr and the reducing power of the sugar was measured with time by the DNS method. [Bmim]Cl: solid line, ChoAc: dashed line. 1 time wash: round, 2 times wash: square, 3 times wash: triangle. The error bars in the figure represent the standard deviation (SD).

To further increase sugar production, we examined the effects of increasing the amount of enzyme, adding enzyme multiple times, and prolonging the reaction time. Based on the results of the experiments conducted thus far, Avicel that was treated with 10% [Bmim]Cl and then washed twice with 50 mM sodium acetate buffer (pH 5.5) was selected as the substrate for subsequent experiments. As shown in Fig. 5, adding two or three times the amount of enzyme (0.076 or 0.114U) increased the amount of sugar produced compared to adding one time the amount of enzyme (0.038U). However, there was not so much difference between adding 2 and 3 times the amount of enzyme. Therefore, we decided to use twice the amount of enzyme (0.076U). We compared the amount of sugar produced when enzymes were added once (0 hr) versus four times (0, 8, 24, and 32 hr) by extending the reaction period up to 48 hours (Fig. 6). In both cases, prolonging the reaction time increased the amount of sugar formation, but adding the enzyme multiple times did not increase the amount formed. The formation rate decreased with increasing reaction time.

**Figure 5.**
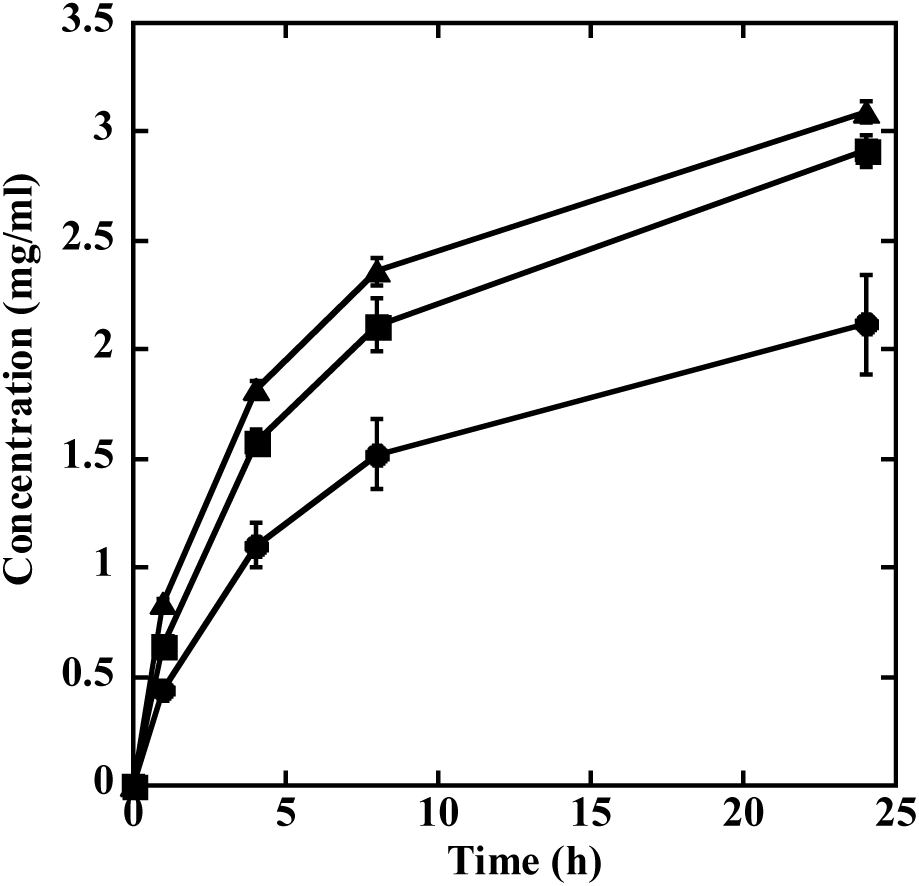
Effect of increasing the amount of enzyme. The 10 mg regenerated Avicel treated with 10% [Bmim]Cl was washed with twice 50 mM sodium acetate buffer (pH 5.5). The resulting pellet was resuspended in 1 mL of 50 mM sodium acetate (pH 5.5) and added CBH HmCel6A-3SNP (0.038, 0.076 or 0.114U) at 0 hr. The reaction was carried out at 80°C for 24 hr and the reducing power of sugars was measured over time using the DNS method. 0.038U: circle; 0.076U: square; 0.114U: triangle. The error bars in the figure represent the standard deviation (SD).

**Figure 6.**
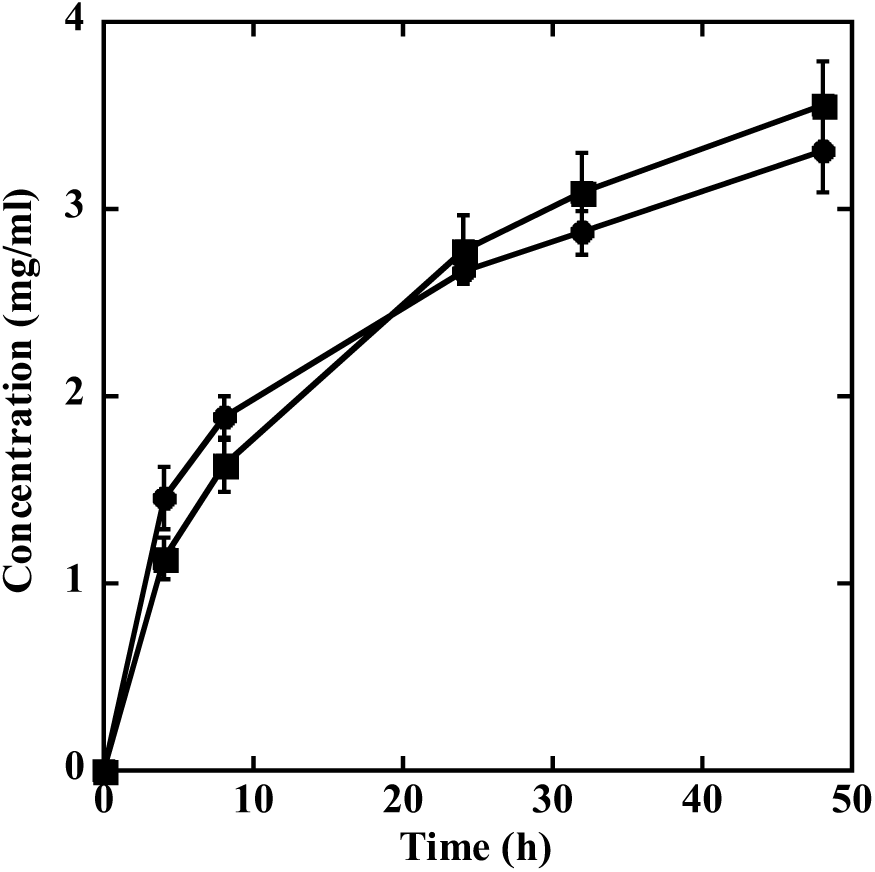
Effect of the number of enzyme additions. The 10 mg regenerated Avicel treated with 10% [Bmim]Cl was washed twice with 50 mM sodium acetate buffer (pH 5.5). The resulting pellet was resuspended in 1 mL of 50 mM sodium acetate (pH 5.5) and added CBH HmCel6A-3SNP (0.076U) at 0, 8, 24 and 32 hr. The reaction was carried out at 80°C for 48 hr and the reducing power of sugars was measured over time using the DNS method. 0 hr only added: circle; 0, 8, 24 and 32 hr added: square. The error bars in the figure represent the standard deviation (SD).

Taken together, these results suggested that the best conditions were to dissolve Avicel with [Bmim]Cl, wash twice with buffer, use 0.076 U/mL of the CBH HmCel6A-3SNP enzyme and allow a 48hr reaction time.

### Identification of enzymatic reaction products

The products from the 48 hr reaction were identified using high-performance liquid chromatography (HPLC) (Table 1). For comparison, the products from commercial cellulases that were reacted under similar conditions were also identified. Under the optimal conditions of this experiment, when CBH HmCel6A-3SNP was used, most of the product was cellobiose, and glucose accounted for 4% of cellobiose. Combining CBH HmCel6A-3SNP with ionic liquid-treated Avicel achieved 37.4% saccharification, and the cellobiose purity in the produced sugars was very high, at about 96%. Using commercial celluloses derived from *T. reesei* and *T. viride*, both enzymes produced nearly the same amount of cellobiose as HmCel6A-3SNP. However, the primary products were both glucose with yields approximately 1.7 times higher than those of cellobiose. The *A. niger* enzyme produced only small amount of cellobiose, with glucose being the dominant component.

**Table 1.** Analysis of products by HPLC.

| Origin | Cellobiose (mg/ml) | Glucose (mg/ml) | Ratio (Glucose/Cellobiose) | Recovery (%) |
| --- | --- | --- | --- | --- |
| HmCel6A-3SNP | 3.595 ± 0.136 | 0.144 ± 0.017 | 0.040 | 37.4 |
| *T. reesei* | 3.304 ± 0.464 | 5.782 ± 0.368 | 1.750 | 90.9 |
| *T. viride* | 2.974 ± 0.380 | 4.840 ± 0.271 | 1.627 | 78.1 |
| *A. niger* | 0.088 ± 0.013 | 7.260 ± 0.397 | 82.50 | 73.5 |

The 10 mg Avicel treated with 10% [Bmim]Cl was washed twice with 50 mM sodium acetate buffer (pH 5.5). The resulting pellet was resuspended in 1 mL of 50 mM sodium acetate (pH 5.5) and then 0.076 U of each enzyme was added. The reaction was carried out at 80°C for CBH HmCel6A-3SNP and at 40°C for the other enzymes. After 48 hr, the concentrations of cellobiose and glucose in the solution were determined by HPLC. The errors indicate the standard deviation (SD) of triplicate samples from the same reaction.

### Structural changes of cellulose by ionic liquid treatment

To examine the degree of destruction of the higher order structure of cellulose by ionic liquids, we observed the birefringence of cellulose using a microscope. Fig. 7 shows the birefringence imaging data of Avicel after being treated with [Bmim]Cl or ChoAc at 100°C for 30 min. Since water treatment does not dissolve cellulose, the large agglomerates with strong contrast in the bright-field image are considered to be the original appearance of Avicel: thick and bulky agglomerates (Fig. 7A). In the brightness image, regions of low retardation with strong contrast were presumed to be formed from multiple crystal with different orientations stacked over and over, which cancels out birefringence. Conversely, regions of high retardation with weak contrast in the bright-field image may represent a crystal clump with an approximately uniaxial orientation. Phosphoric acid-swollen cellulose (PASC), which is susceptible to degradation by cellulase, exhibited reduced birefringence given no molecular orientation in the optical microscopic scale (Fig. 7H).

**Figure 7.**
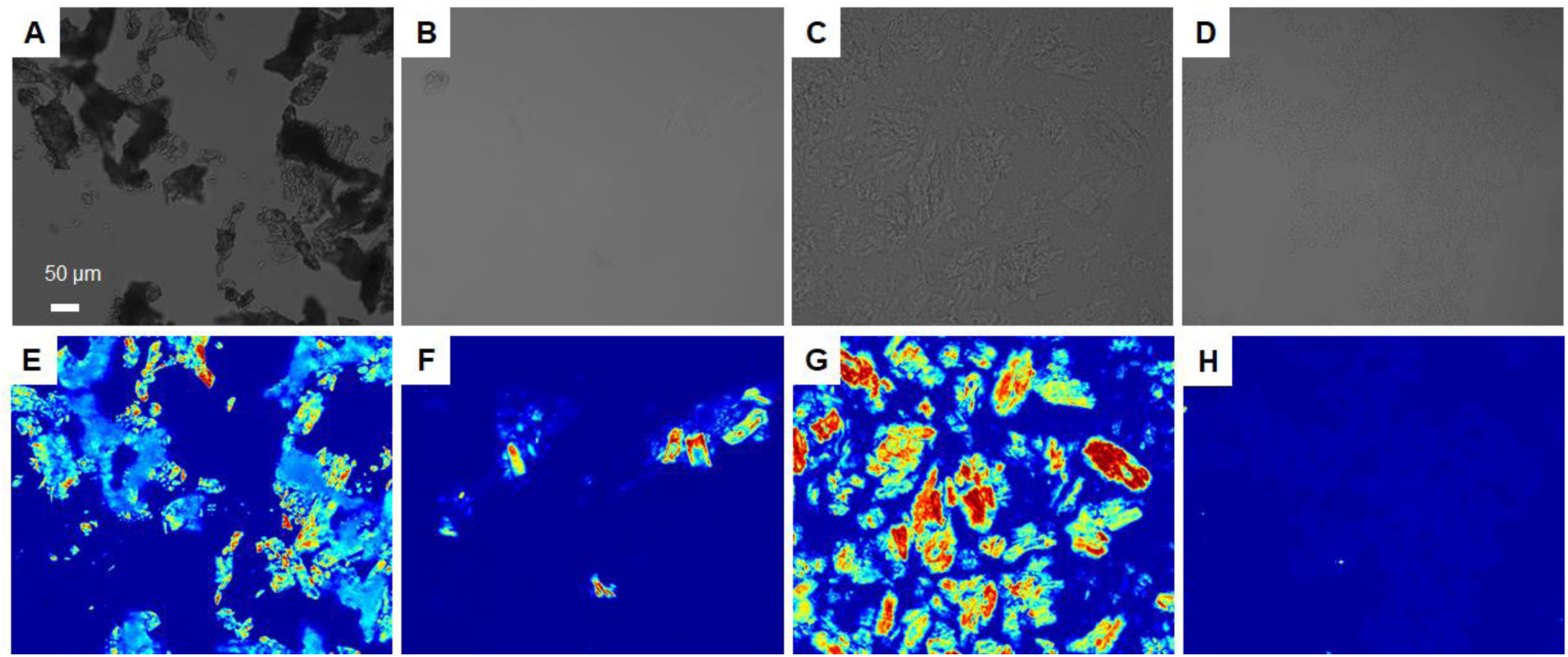
Birefringence imaging of Avicel treated with water, [Bmim]Cl, ChoAc and PASC at 100°C for 30 min. The retardation values are shown in a color-coded scale ranging from blue (0 nm) to red (greater than 130 nm). A-D are the bright-field image and E-H are the birefringence image. A, E: Water treatment; B, F: [Bmim]Cl treatment; C, G: ChoAc treatment; D, H: PASC

Meanwhile, the contrast of Avicel treated with [Bmim]Cl or ChoAc was clearly weaker than that of Avicel treated with water in the bright-field images (Fig. 7B and C), indicating the disintegration of the cellulose aggregation. This was considered to be due to the dissolution of microcrystalline cellulose, but it could not be ruled out that the high refractive index of the ionic liquid. After [Bmim]Cl treatment, however, the regions with high retardation values became less significant in the observation. In contrast, after ChoAc treatment, cellulose clumps with high retardation values were observed, unlike before treatment. These observations confirmed that the aggregation present in the untreated cellulose were disintegrated, indicating that the cellulose was at least partially dissolved in the ionic liquids (Fig. 7F, G). Considering the cellulose bundles with exceptionally high retardation (Fig. 7G), it seemed that ChoAc solely disintegrated the cellulose bundles within a large particle, rather than dissolving the cellulose itself. However, this possibility was ruled out by an interesting observation regarding Avicel treated with ChoAc (Fig. 8). After the heating treatment, the ionic liquid-treated Avicel sample was transferred directly to a slide glass and observed immediately at room temperature. Initially, we observed primarily 50 μm or larger cellulose aggregation with relatively high retardation and, a background with almost zero retardation, as discussed in Fig. 7G. After several minutes, however, the regions of very low retardation disappeared and highly birefringent regions spread throughout the sample, as shown in Figs. 8C and 8D. Fig. 8C is a snapshot of this transition. This is probably due to cellulose regenerating from a dissolved to a solid state, suggesting that dissolved cellulose molecular chains exist in the region of low retardation in Fig. 8C.

**Figure 8.**
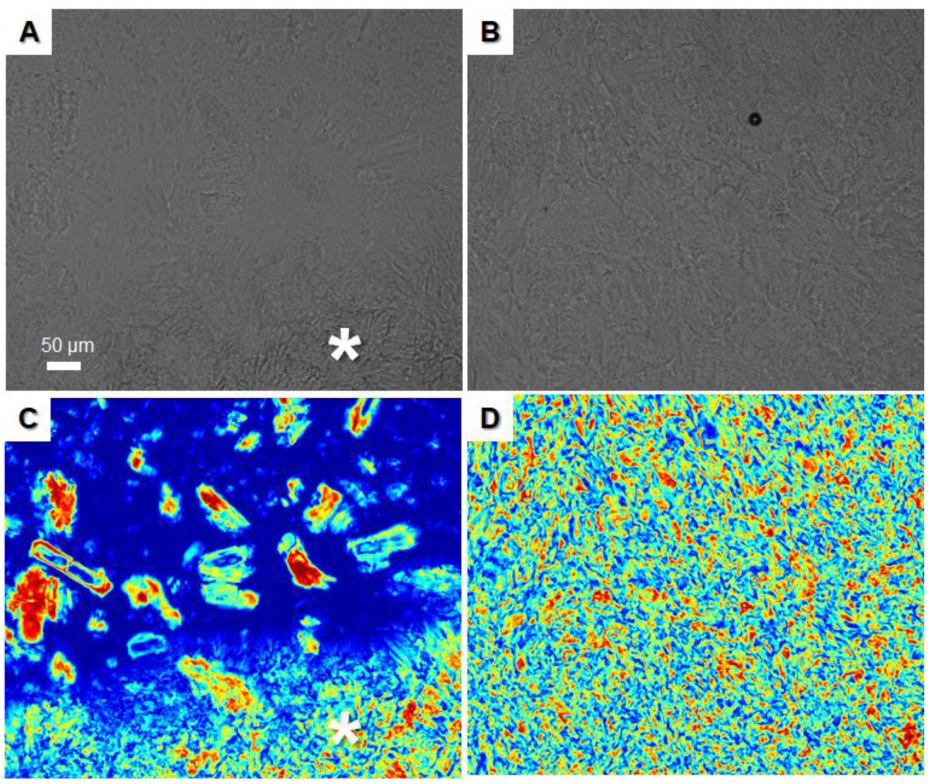
Birefringence imaging of Avicel treated with ChoAc at 100°C for 30 min. The retardation values are shown in a color-coded scale ranging from blue (0 nm) to red (greater than 130 nm). A and B are bright-field images, and C and D are birefringence images. A and C show cellulose that is still in a dissolved state, as well as the part that is solidifying. The area indicated by the asterisk is the solidified area, and the upper part shows the dissolved area of cellulose before solidification. This asterisk area eventually covered the entire visual field during observation, until after a few minutes, the entire area was marked D.

### *in situ* birefringence imaging

The decay of birefringence during treatment was visualized by *in situ* observation of cellulose in [Bmim]Cl or ChoAc using a birefringence detector (Fig. 9). The clear decrease in birefringence of the cellulose when [Bmim]Cl was used indicates that molecular orientation was lost during cellulose dissolution (Fig. 9A-F). This dissolution proceeded gradually up to 100°C and was nearly complete after 90 min at 100°C. Therefore, higher temperatures and longer incubation period would promote the dissolution of Avicel in [Bmim]Cl. Conversely, after 90 min incubation at 100°C, no loss of birefringence was observed when ChoAc was used (Fig. 9G-L), indicating that ChoAc did not dissolve the highly crystalline cellulose Avicel at 100°C. However, retardation increased slightly after 90 min treatment with ChoAc (Fig. 9L). This observation may reflect an improvement in the orientation of the cellulose molecules within the solid cellulose, yielding a higher retardation. It is possible that incubating in ChoAc at 100°C enhances the rearrangement of cellulose molecules in the crystalline part and/or the reabsorption of dissolved cellulose molecules onto the crystal surface. This hypothesis requires testing, but given the lack of significant structural changes observed by light microscopy, it remains the most plausible explanation.

**Figure 9.**
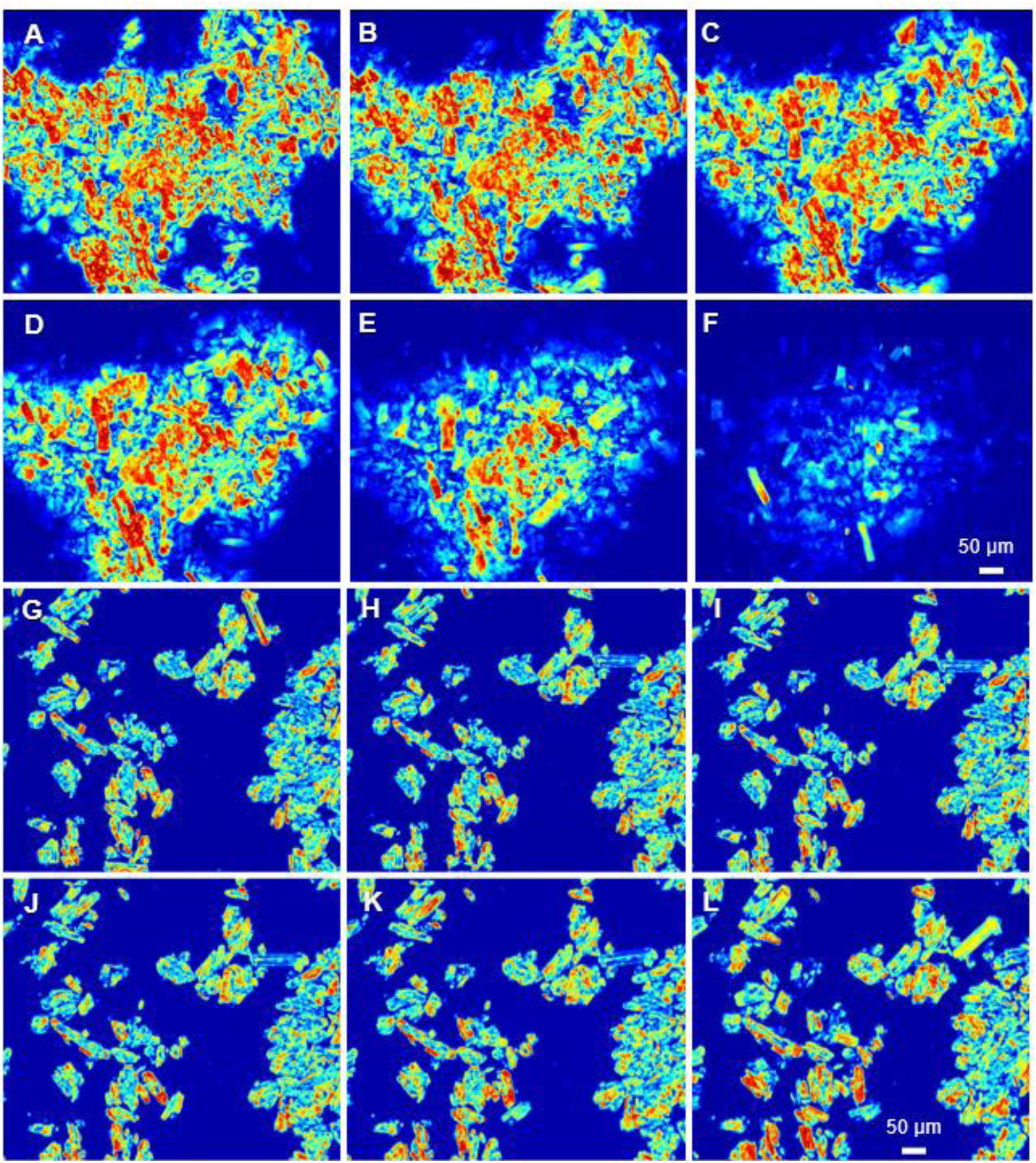
Time-lapse imaging of cellulose in an ionic liquid during heating using a birefringence detector. Retardation values are shown in a color-coded scale ranging from blue (0 nm) to red (greater than 130 nm). (A-F) Changes of Avicel in [Bmim]Cl: (A) 70°C; (B) 80°C; (C) 90°C; (D) 100°C; (E) 100°C for 30 min.; (F) 100°C for 90 min. For A-D, the temperature was maintained for 2 min after jumping to the next set temperature before recording. (G-L) Changes of Avicel in ChoAc: (G) 70°C; (H) 80°C; (I) 90°C; (J) 100°C, (K) 100°C for 30 min; (L) 100°C for 90 min. For G-J, the temperature was maintained for 2 min after jumping to the next set temperature before recording.

### SEM observation

Structural changes in the surface morphology of cellulose were also confirmed by scanning electron microscopy (Fig. 10). Bulky features were observed in all samples except for PASC. This may explain in part why PASC is easily degraded by many cellulases. In other words, these enzymes can more easily access PASC than bulky cellulose samples. In Avicel, large particles of cellulose aggregation (ca. 10-100 μm) were frequently observed. he cellulose microfibrils within these particles were tightly packed, and there were no apparent interfibrillar spaces (Fig. 10A and B). Since these dispersed particles were not observed after treatment with either ionic liquid (Fig. 10C and E), these particles might have fused due to the ionic liquid treatment. At higher magnification, the fiber morphologies and interfibrillar voids were clearly visible, indicating that both ionic liquids significantly interfered with the structure of Avicel (Fig. 10D and F). At low magnification, the difference between [Bmim]Cl- and ChoAc-treated Avicel was evident: thin fabric structures were seen in the former, whereas denser aggregates (>100 μm) were mainly observed in the latter.

**Figure 10.**
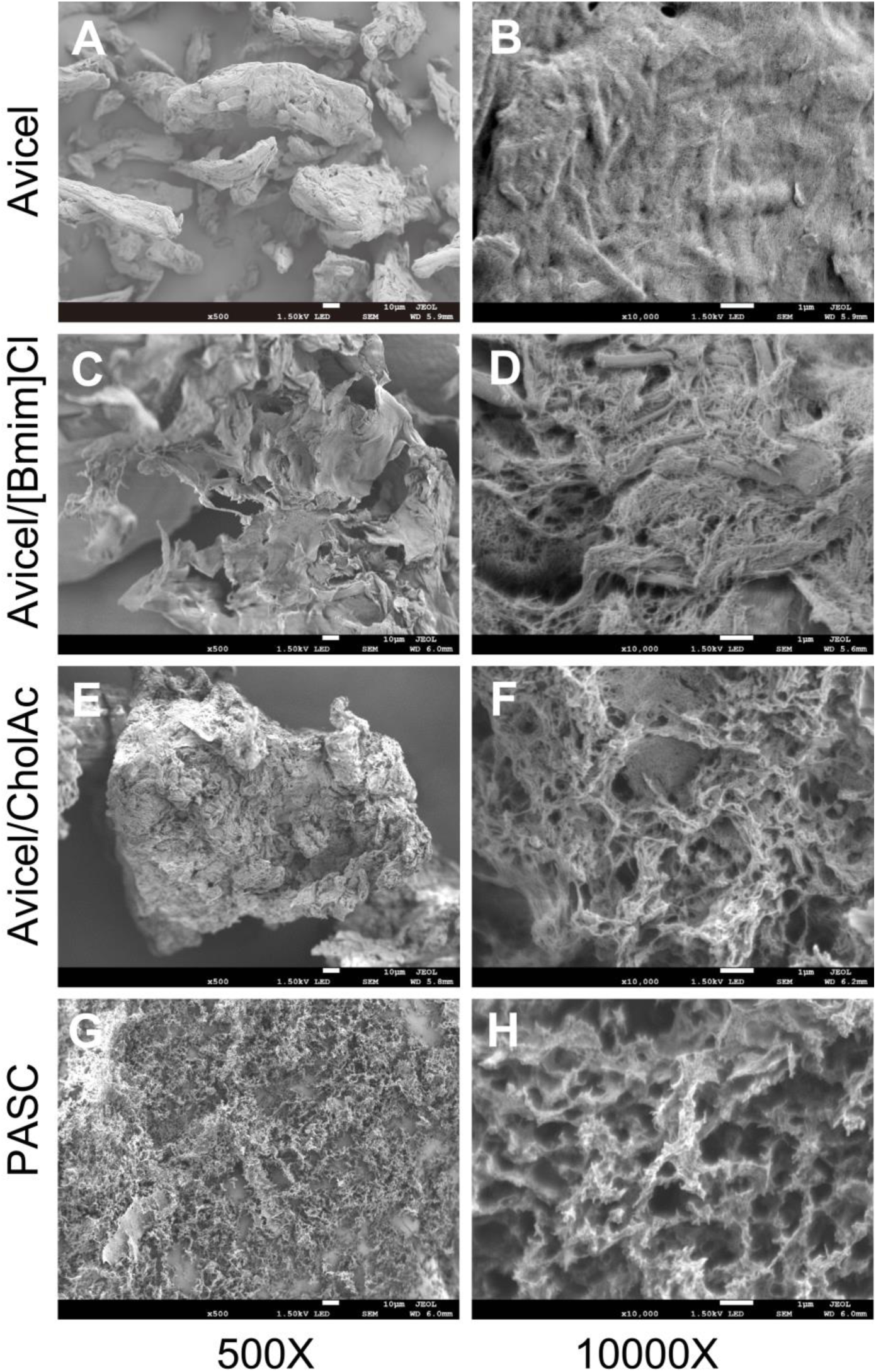
Scanning electron micrographs of ionic liquid-treated cellulose and control samples (Avicel and PASC). Left: low magnification (500 fold), Right: high magnification (10,000 fold)

### FTIR spectroscopy

Attenuated total reflection Fourier transform infrared spectroscopy (ATR-FTIR) was used to analyze the internal structure of cellulose treated with ionic liquids. Fig. 11A shows the full spectral range, and Fig. 11B shows a close-up of the OH-stretching region between 3700 and 3000 cm^-1^. The figures display the spectra of original Avicel, [Bmim]Cl-treated Avicel, ChoAc-treated Avicel, and PASC. A typical spectrum clearly indicated that both [Bmim]Cl- and ChoAc-treated Avicel were cellulose II due to the specific features of cellulose II in the OH stretching region (3440 and 3485 cm^-1^, indicated by dotted bars in Fig. 11B). Avicel treated with ChoAc did not clearly demonstrate the dissolution process in birefringence imaging (Fig. 9). However, ATR-FTIR revealed that the ChoAc treatment was also converted Avicel into cellulose II crystals. Notably, the IR spectra of these ionic liquid-treated Avicel samples have fewer sharp features compared to PASC, indicating lower crystallinity. PASC is well-known as a typical cellulose with low crystallinity. The study demonstrated that the ion-liquid-treated Avicel samples have an even lower crystallinity than PASC. This likely explains why they are more digestible by CBH HmCel6A-3SNP than PASC.

**Figure 11.**
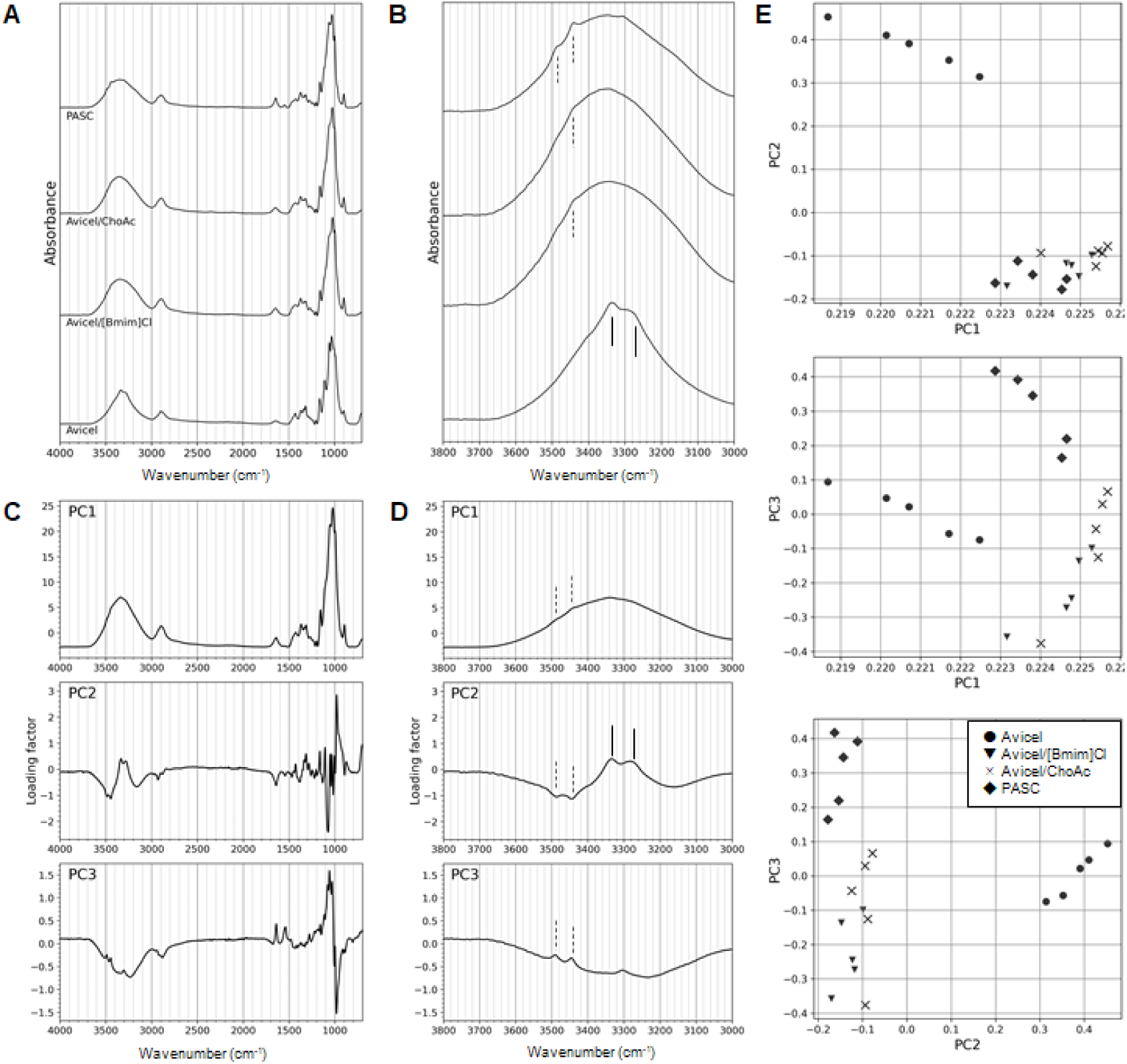
FTIR spectra and PCA analysis of cellulose samples. A and B: Typical spectra from Avicel, [Bmim]Cl-treated Avicel, ChoAc-treated Avicel, and PASC. Panel B is a close-up of the OH-stretching region between 3700 to 3000 cm^-1^. The solid and dashed vertical bars in the spectrum represent spectral features specific to cellulose I and cellulose II, respectively. (C and D) The loadings of the first three principal components obtained from the PCA analysis. Panel D is a close-up of the OH-stretching region between 3700 to 3000 cm^-1^. The solid and dashed vertical bars indicate spectral features unique to cellulose I and cellulose II, respectively. The dotted vertical bar is present at about 3300 cm^-1^ between the peaks at 3285 and 3355 cm^-1^ characteristic of cellulose I. The contribution ratios of PC1, PC2, PC3 and PC4 are 0.97146, 0.018941, 0.007543 and 0.001082, respectively. (E) Score plot of two principal components from PCA analysis: PC1-PC2, PC2-PC3, and PC1-PC3.

PCA of the IR spectra was then performed to search for hidden spectral features that might explain the efficiency of CBH HmCel6A-3SNP reaction (Fig. 11C, D and E). The loading of the first principal component (PC1) was very similar to that of the original spectrum. This allows us to interpretate PC1 as representing characteristic of either cellulose I or II crystals, given its 97% contribution to the dimension-reduced model. The other principal components are likely to explain the differences between the samples. The PC2 and PC3 loading factors, as indicated by the OH stretch region, appear to meaningfully represent the difference between the negative contribution of cellulose II-specific features and the positive contribution of cellulose I-specific features. In particular, PC2 effectively separates the sample groups (cellulose I and cellulose II) between Avicel and others, as shown in Fig. 11D.

Given the reasonable grouping, the PCA results were further assessed to find differences between [Bmim]Cl- and ChoAc-treated Avicel. While the categorization was not as clear as along PC2, PC3 appears to represent the difference between PASC and these ionic liquid-treated Avicel samples. Classification along PC3 ranks the ionic liquid-treated cellulose samples from PASC, and ChoAc-treated Avicel was closer to PASC rather than to [Bmim]Cl-treated Avicel. This trend is consistent with the digestibility of the cellulose samples by CBH HmCel6A-3SNP in this study: [Bmim]Cl-treated Avicel> ChoAc-treated Avicel>PASC. Thus, the PC3 may encode the structural factor responsible for the enzymatic digestibility of cellulose by the CBH.

## Discussion

Here we examined the production of cellobiose from regenerated cellulose with ionic liquids by the action of the highly thermostable CBH HmCel6A-3SNP at 80°C. As mentioned in the introduction, previous studies using thermostable and ionic liquid-tolerant CBH enzymes have only reached a maximum reaction temperature of 65°C or lower. Therefore, this study significantly increased the reaction temperature. The CBH used in this study retained its activity at lower concentrations of [Bmim]Cl or ChoAc ionic liquid solution when PASC was used as the substrate (Fig. 1). Although the inhibitory effects of [Bmim]Cl on the CBH activity were stronger than those of ChoAc (Fig. 1), the time course experiment revealed that the highest amount of cellobiose production was achieved using the ionic liquid-treated Avicel as the substrate in the 10% [Bmim]Cl solution (Fig. 2). Conversely, cellobiose production was lowest in the 20% [Bmim]Cl solution and was comparable to the production level from native Avicel. This is considered to be due to the stronger enzyme inhibitory effect in 20% [Bmim]Cl solution compared to 10% solution (Fig. 1). Both [Bmim]Cl- and ChoAc-treated Avicel showed improved cellobiose productivity when washed with the acetate buffer, the yield was higher with the [Bmim]Cl treatment. (Fig. 3). For both ionic liquids, the first washing cycle proved most effective.

As shown in Fig. 2, it is noteworthy that the initial rates of cellobiose production were similar and within the margin of error for most reactions involving ionic liquid-treated Avicel substrates up to the 1 hr time point (excluding 20% [Bmim]Cl-treated and native Avicel). However, the production rates decreased differentially thereafter. This may suggest that the readily accessible substrates that are more easily hydrolyzed are nearly consumed by the amount of enzyme added at the beginning of the reaction after about 1 hr. The remaining substrates may then react at different rates depending on their features. This hypothesis is supported by the experiment in which more enzyme was added at the beginning of the reaction to the [Bmim]Cl-treated regenerated cellulose, resulting in a cellobiose production curves approaching saturation (Fig. 5). The experiment involving the addition of an equal amount of enzyme to the reaction at 8, 24, and 32 hr showed similar reaction features to those without such additions (Fig. 6), which also supports the hypothesis. With these enzyme additions, the amount of cellobiose production was nearly reached saturation at 36% of the total amount of cellulose, while commercially available cellulase cocktails containing endoglucanases and beta-glucosidases exhibited higher sugars recovery rates of over 70% from the regenerated cellulose (Table 1). Taken together, these results suggest that the ionic liquid-regenerated cellulose substrates were partially accessible to the CBH used in this study. The differentiated accessibility of these regenerated cellulose substrates to the enzyme suggests that their structural features differ after regeneration. Therefore, we investigated the structural features, as shown in Figs. 7 to 11.

It has been previously reported that conversion of the crystal structure of cellulose from the native type I crystal to another crystal structure increases its enzymatic saccharification property (Igarashi *et al*. 2007, Kobayashi *et al*. 2012; Wada *et al*. 2010). Consistent with these reports, the ionic liquid-treated cellulose converted from cellulose I to cellulose II tends to show higher enzymatic saccharification properties than the native Avicel. In addition, the lower crystallinity of [Bmim]Cl-treated cellulose compared to PASC, as demonstrated by FTIR, suggests that the degree of the molecular chain disorder is closely related to enzymatic saccharification properties. However, the fact that the enzymatic saccharification of the ChoAc-treated cellulose was not as effective as that of the [Bmim]Cl-treated cellulose(Figs. 2-4). Additionally, FTIR analysis revealed that the crystallinity of the ChoAc-treated cellulose was as low as that of [Bmim]Cl-treated cellulose. This suggests that the enzymatic saccharification cannot be explained by the crystallinity alone. PCA was used to verify whether there were any feature values in the FTIR spectrum that could clearly explain the enzymatic saccharification properties. PC3 tended to correlate with the saccharification property. Therefore, the loading factor of PC3 may reveal spectral features related to enzymatic saccharification. The [Bmim]Cl-treated cellulose exhibited the highest potential for enzymatic saccharification. Some characteristics can be interpreted as indicating an increase in saccharification potential with weaker crystallinity. For example, weak absorption was observed at 3440 and 3485 cm^-1^ in the OH-stretching region, which is characteristic of cellulose II. However, other characteristics could not be easily interpreted, such as low absorption at 1040/1060 cm^-1^ and high absorption at 980 cm^-1^ in the finger-print region, which require further verification.

On the other hand, the relatively large-scale structures observed by the birefringence camera and SEM were crucial in explaining the enzymatic digestibility by the CBH used in this study. Avicel was dissolved by treatment with ionic liquids and reaggregated by washing with the buffer solution. SEM observation showed that the degree of aggregation was weak for [Bmim]Cl-treated cellulose, but relatively strong for ChoAc-treated cellulose.

In the birefringence image, a clear, solid structure was observed in the cellulose treated with [Bmim]Cl. However, in the case of ChoAc treatment, a solid structure was observed without a clear dissolution process, although it was presumed that some molecules had been dissolved (Fig. 9). In both cases, FTIR revealed significant structural changes from cellulose I to cellulose II, suggesting that the [Bmim]Cl-treated cellulose underwent a process similar to the formation of regenerated cellulose such as rayon. In contrast, the ChoAc-treated cellulose underwent a solid structural transformation, comparable to mercerization using a concentrated alkaline solution. SEM observation showed that the [Bmim]Cl-treated cellulose formed a layered structure, while the ChoAc-treated cellulose exhibited a bulky and relatively densely packed structure. These results reflect the preparation process of the cellulose samples and suggest that enzyme accessibility impacts enzymatic saccharification.

Thus, the present study successfully demonstrated the effect of structural changes in cellulose caused by ionic liquid pretreatment on enzymatic saccharification. Although accumulating such knowledge is challenging, it is important for developing future guidelines that will improve the efficiency of enzymatic saccharification.

## Acknowledgments

This work was supported by the Honda R&D Co., Ltd. SEM observation was performed with a collaborative research facility CAN-DO in RISH, Kyoto University.

